# Polygenic prediction and gene regulation networks

**DOI:** 10.1101/2024.05.07.592928

**Authors:** Juan F Poyatos

## Abstract

Exploring the degree to which phenotypic variation, influenced by intrinsic nonlinear biological mechanisms, can be accurately captured using statistical methods is essential for advancing our comprehension of complex biological systems and predicting their functionality. Here, we examine this issue by combining a computational model of gene regulation networks with a linear additive prediction model, akin to polygenic scores utilized in genetic analyses. Inspired by the variational framework of quantitative genetics, we create a population of individual networks possessing identical topology yet showcasing diversity in regulatory strengths. By discerning which regulatory connections determine the prediction of phenotypes, we contextualize our findings within the framework of core and peripheral causal determinants, as proposed by the omnigenic model of complex traits. We establish connections between our results and concepts such as global sensitivity and local stability in dynamical systems, alongside the notion of sloppy parameters in biological models. Furthermore, we explore the implications of our investigation for the broader discourse surrounding the role of epistatic interactions in the prediction of complex phenotypes.

**Author Summary:** This research delves into how well statistical methods can capture phenotypic variation influenced by nonlinear biological mechanisms. The study combines a computational model of gene regulation networks with a linear additive prediction model, similar to polygenic scores used in genetic analysis. By creating a population of individual networks with identical topology but varying regulatory strengths, the research identifies key regulatory connections that predict phenotypes. The findings are framed within the omnigenic model of complex traits, distinguishing core and peripheral causal determinants. The study also links its results to concepts like global sensitivity and local stability in dynamical systems, as well as sloppy parameters in biological models. Additionally, it examines the implications for understanding the role of epistatic interactions in predicting complex phenotypes. This work enhances our understanding of complex biological systems and their functionality.

## Introduction

One of the most challenging questions in biology revolves around anticipating phenotypes [1]. From a fundamental perspective, the ability to predict phenotypes, particularly complex ones, sheds light on the coordination among genetic, cellular, and organismal levels [2]. It also elucidates the emergence of robustness and the consequences of its failure, leading to malfunction [3]. With directed application, this knowledge empowers us to intervene more effectively in biology, redirecting disease trajectories, and engineering novel systems with desired properties [4].

However, two complementary research avenues provide seemingly contrasting viewpoints on the accomplishment of this challenge. On one hand, quantitative genetics has successfully developed statistical frameworks that integrate genetic and phenotypic variation to predict phenotypes. These frameworks, which are generally additive, have been effectively applied in agriculture and livestock [5] and also extended into human purposes with the development of polygenic scores (PGSs, see glossary box), which quantify the genetic predisposition of *individuals* to specific traits or diseases [6–8]. On the other hand, molecular genetics is increasingly delineating numerous mechanisms contributing to complex traits, emphasizing the many nonlinear processes involved [9]. The presence of nonlinearities at the mechanistic level seems to present a potential limitation to prediction.

These apparent contradictions call for a reassessment of our approach to this problem. One aspect of this revision involves a fundamental discussion on the limitations of variance decomposition in quantitative genetics [10]. This decomposition is not inherently linked to the mechanistic aspects of phenotype generation or gene action [11]. Indeed, the existence of interactions between genes, referred to as biological or functional epistasis, does not necessarily correspond solely to the nonadditive component of variance [12]; it may, in fact, contribute to the additive component [13–16]. Consequently, it is not entirely clear how to reconcile these two aspects or if it is even possible [17].

Moreover, the notion that tools like PGSs would rest on relatively few and strong-effect determinants has been reevaluated in humans, recognizing the abundance of loci with small effect sizes related to complex traits. This is the omnigenic model [18, 19], which defines the genetic determinants of phenotypes in terms of causality, identifying “core” genes with direct effects on the phenotype and “peripheral” genes that influence phenotypes indirectly. This reformulation underscores the perspective of phenotypes as emergent properties of interacting components within networks, supporting the systemic interpretation of biology advanced in recent years [19–21]. Nonetheless, arguments regarding the relevance of biological networks in this context are often qualitative and somewhat incomplete.

In this work, we aim to provide a formalized examination of this view. To this end, we utilize a mathematical framework that simulates the action of a gene regulation network constituted by random regulatory relationships between genes with a given strength [22, 23]. The analysis is not intended to thoroughly explore all possible regulatory topologies in specific contexts but rather aims to provide an initial understanding of how phenotypes produced by gene networks can be predicted using statistical tools. Thus, we use this simplistic *generative* model to emulate a variational situation in which a population of networks exhibits genetic and phenotypic variation relative to a reference structure. Genetic variation is represented by the differences in regulatory weights around the values of the reference network, while phenotypic variation is associated with the resulting steady-state expressions of the genes. Importantly, the data generated in this manner enables us to calculate a PGS following the approach of quantitative genetics with the added advantage of examining the mechanism generating the phenotype, which is the regulatory network.

Using this framework, we examine how the PGS is determined by the intrinsically nonlinear regulatory connections and identify any limitations on its prediction accuracy. To achieve this, we produce a variety of reference networks and investigate how the range of linear functioning, the levels of genetic variation, or the occurrence of minor changes in network structure, influence the statistical predictions. As PGSs are also dependent on the distribution of genetic variation used to their characterization, we will also quantify how well these methods predict the phenotype of individuals drawn from a different distribution, a problem commonly referred to as “transferability”, e.g., [24]. Finally, we delineate core and peripheral causal elements inspired by the omnigenic framework and their relationship with intrinsic features of the mechanisms leading to the phenotype. More broadly, our study provides insights into the predominant role of strictly additive models in describing the relationship between genotype and phenotype.

### Glossary

**Individual prediction:** the prediction of phenotypic traits at the individual level, taking into account the unique genetic makeup and environmental factors of the individual.

**Statistical prediction:** mathematical models that use statistical methods to predict or explain certain outcomes using population data.

**Polygenic score (PGS):** an additive statistical model that is typically calculated based on the presence of multiple alleles, each contributing to a particular trait or disease risk.

**Transferability:** it refers to how effectively a given PGS, derived under specific conditions (e.g., genetic variation within a population), can predict traits under different conditions.

**Variance decomposition:** a technique developed in quantitative genetics that decomposes the phenotypic variance in a population as a sum of genetic and environmental variation.

**Genetic architecture:** it refers to the number, frequency, and effect sizes of genes and their interactions, that collectively influence the expression of a trait.

**Epistasis:** interaction between genes that can refer to statistical or physiological aspects.

**Gene regulation network (GRN):** a network in which the nodes correspond to a set of genes, which typically encode a transcription factor, and the edges indicate regulatory interactions between them.

**Genetic and phenotypic variation in GRNs:** genetic variation refers to differences in regulatory strengths between genes across individual GRNs within a population, while phenotypic variation is the resulting change in the average steady-state expression of the GRN’s genes.

**Omnigenic model:** a model that suggests that most genes contribute to complex traits, with “core” genes having direct effects and “peripheral” genes influencing traits indirectly through GRNs.

**Local stability analysis:** a method that examines the stability of a dynamical system’s equilibrium by analyzing small perturbations around that equilibrium.

**Global sensitivity analysis:** a procedure that assesses how variations in model inputs influence the outputs across the entire input space. It produces Sobol’s additive and nonadditive indices.

**Network dynamical nonlinearity:** describes the nonlinear behavior of a system arising from interactions within a network with the use of different mathematical methods.

## Results

### A polygenic score that predicts the phenotype of a regulatory network

We first analyze a population consisting of individual gene regulation networks, where nodes represent genes and edges denote the regulatory interactions between them. In this population, the network structure is fixed, but there is variation in the interaction strengths, which deviate from a set of defined values (constituting the *reference network*, e.g., Fig. 1A, Methods). This variation in strength, or weight, can reflect mutations in DNA regulatory regions.

**Figure 1.**
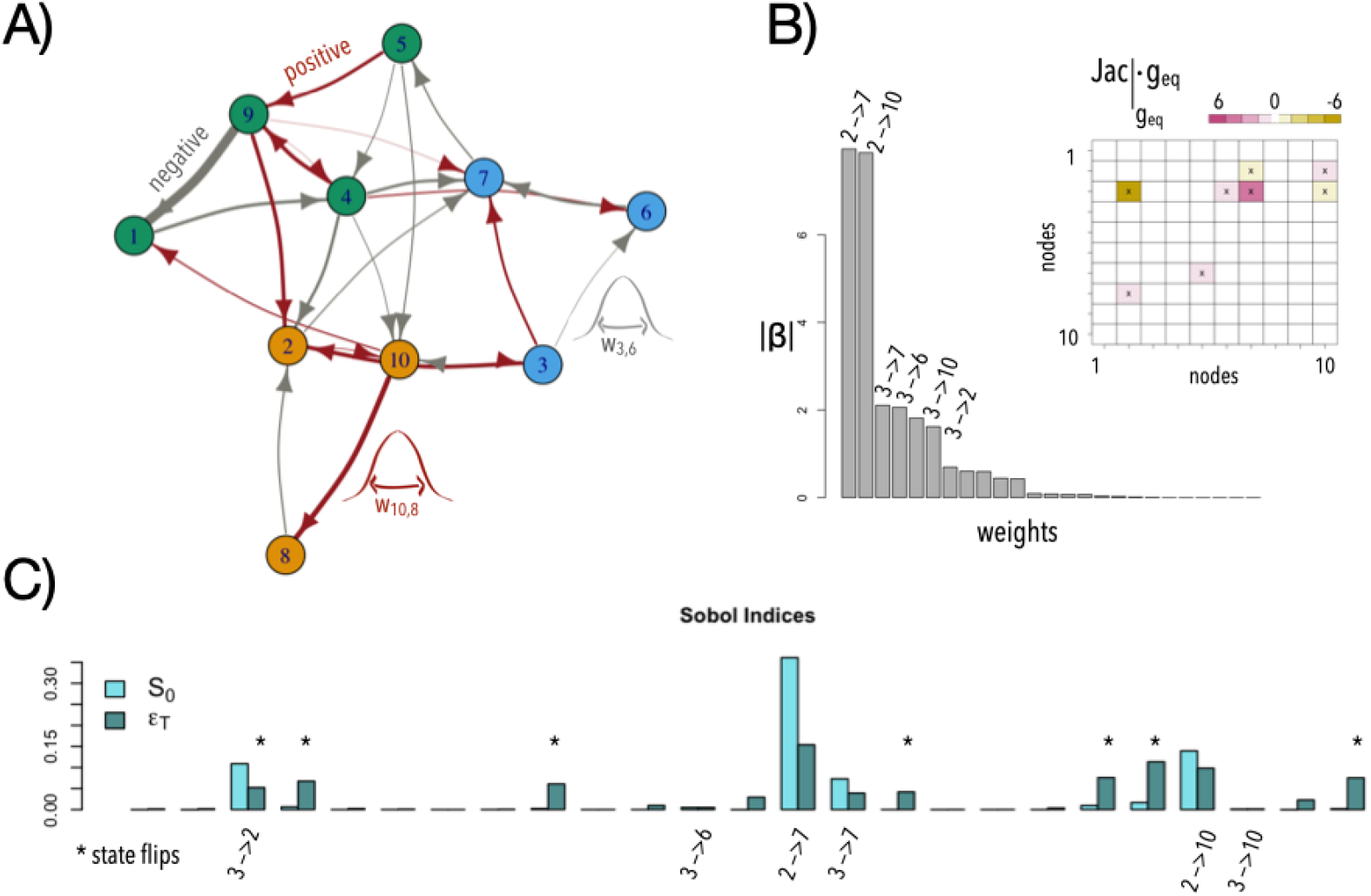
PGS, local stability and global sensitivity analysis of a model network. **A)** Example of a *reference* network with *N* = 10 genes. We develop a variational framework in which each network in the population exhibits the same structure, but its weights are sampled from normal distributions centered around the weights of the reference network, with variations around 10% of these values (sd = 0.1, two example distributions are shown). Red and gray edges represent positive and negative interactions, respectively. Nodes are colored here following a simulated annealing procedure to identify communities in a network. **B)** Absolute value of effect size, |β|, for each regulatory connection in the PGS obtained. Top values are associated to their respective edges. Inset. The product of the Jacobian matrix at equilibrium and the steady-state gene expression of the reference network predicts the weights of the top contributors in the PGS. Color highlights the strength of the values. **C)** Sobol indices where 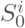 describe the additive contribution and 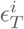 the nonadditive contribution of each edge *i* to the variation of the phenotype. Asterisks indicate those weights that contribute to state flips when subjected to variation (see main text). Here *i* → *j* represents the regulation of *i* on *j* with weight *w*_*i,j*_. Time to reach equilibrium (“path length”) of model network = 18 (time steps), PGS *R*^2^ = 0.69.

For each network, we calculate the average of the steady-state expression values of its genes, which serves as the phenotype. The steady state is obtained by iteratively multiplying the regulatory matrix starting from a particular initial expression-level vector, with each element of the resulting vector at each iteration being transformed by a sigmoidal function, thereby mirroring the behavior of actual gene regulation networks (Methods) [22, 23].

We use this comprehensive dataset, incorporating genetic variation (in weights) and phenotypic variation (in average steady-state expression), to derive a multidimensional PGS (Methods). In the PGS, each regulatory interaction contributes with an additive effect size (β) as predictor of the phenotype. Figure 1B shows the specific β coefficients (absolute values) associated with the regulatory connections depicted in Fig. 1A. The corresponding PGS predicts ∼70% of the variation in phenotype observed in the population.

### PGS determinants contribute both additively and non-additively to the phenotype

To further investigate the rationale behind the β coefficients in the PGS, we leveraged our knowledge of the mechanistic network generating the phenotypes. Firstly, given the linear and additive nature of the PGS, we applied local stability analysis, which involves assessing how small perturbations in a dynamical system affect its equilibrium (steady) state [25]. To this aim, one can linearize the regulatory interactions to determine which ones predominantly contribute to deviations from the reference phenotypic value (colored squares, inset of Fig. 1B; this is achieved by computing the Jacobian matrix of the dynamical equations for the reference network at the equilibrium, then multiplying it by the gene expression levels at the steady state, Methods). These connections, which cause the network to behave more linearly in that specific region, correspond to those with larger effect sizes in the PGS.

However, local stability analysis does not fully explain the *β*’s as it is limited to slight variations away from the steady state. We thus adopted global sensitivity analysis, a technique that allows us to quantify how variations across the weight parameter space, particularly crucial in nonlinear regulatory networks, lead to phenotypic variation. We can then decompose phenotype variance into contributions from individual weights and from collective sets of weights [26]. Specifically, this method yields first-order 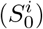 and total-order 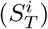 effect indices –Sobol’s indices– for each regulatory connection *i*, where the former quantifies the main additive effect contribution to the variance of the phenotype of each connection in isolation and the latter encompasses both additive and nonadditive effects. From these, we derive the total contribution of nonadditive effects as 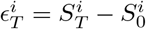.

Figure 1C shows this analysis for the same variational dataset as before. Strong additive contributions (cyan bars in the figure) identify connections with strong *β*^*′*^s, e.g., 2→7, 2→10, etc. These determinants also show nonadditive contributions to the variation in the phenotype. More broadly, nonadditive contributions stress weights whose alteration often results in a significant change in the state of the regulated gene, a state *flip*, inducing a nonlinear effect that percolates throughout the entire network (S1 FigureA). Note also that in this exemplary case the sum of first-order effects is approximately 0.72 (what explains the value of *R*^2^ in the PGS), while the sum of the total indices is 1.58; as these two sums are both different from 1, the influence of any single regulation usually depends on the state of others [27].

### Complex networks typically result in weaker predictability

Given the high coefficient of determination of the PGS for the exemplary network (*R*^2^ = 0.69, Figure 1), it is reasonable to hypothesize that variations in weights lead to proportional changes in the phenotype, a behavior that can also be assessed through the network’s dynamical response, as revealed by the eigenvalues of the Jacobian matrix [25]. For this example, all eigenvalues are much smaller than one, indicating that the system quickly returns to its original phenotype after a quantitative regulatory change, thereby behaving almost linearly near the equilibrium phenotype (Methods).

However, the operation of regulatory networks can transition from fairly linear to strongly nonlinear regimes [28]. Therefore, to systematically examine the boundaries of prediction accuracy, we generated a set of 100 random networks with the same number of nodes and expected connectivity (Methods). We observed instances of both linear and nonlinear behaviors, as defined with the help of stability analysis (S1 FigureBC). These mechanistic features are partially captured by the statistical indices *S*_0_ (the sum of all individual interactions’ additive indexes, 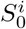, equivalent to *R*^2^ [27]) and ϵ_*T*_ (the sum of all nonadditive indexes, 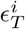), respectively. These parameters determine the location of a network in what we termed the *additive space* (Figure 2).

**Figure 2.**
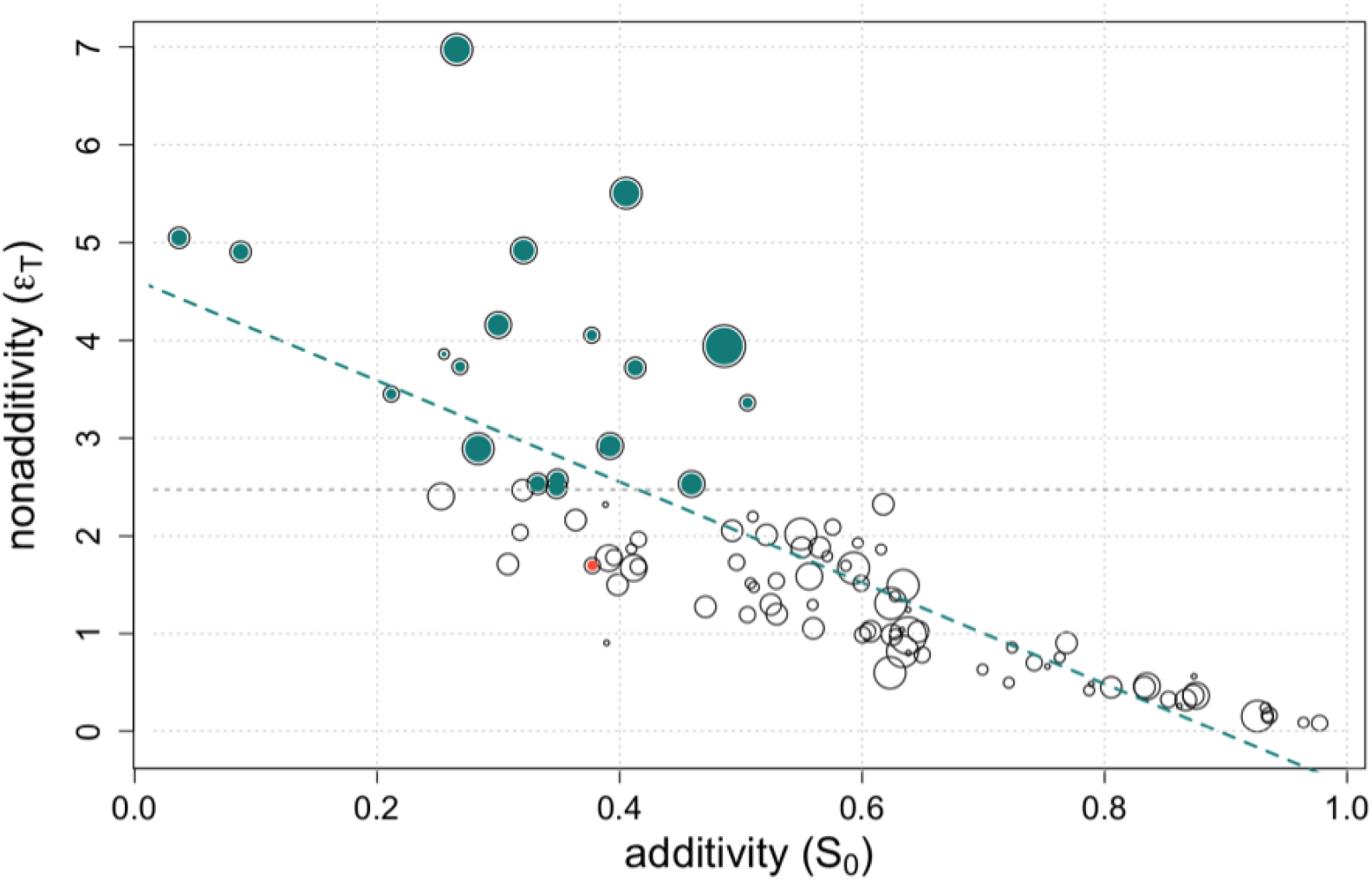
The additivity space of gene regulation networks and the presence of two-component positive feedbacks. We computed the sum of Sobol’s additive (*S*_0_) and nonadditive (ϵ_*T*_) indexes for each network across a dataset consisting of 100 random networks. Dot size represents the number of two-component positive feedbacks found on each network normalized by the median value of the entire population (median = 2). Networks with nonadditivity values beyond the 80th quartile of the population (indicated by the dashed gray line, networks highlighted in green) are enriched in two-component positive feedbacks (observed mean number = 3.3, expected mean number = 2.47, *p*-value = 0.005 after 10000 permutations). The red dot denotes a network with two two-component positive feedbacks, while the green dashed line illustrates the additive-nonadditive effects trade-off.

We then investigated whether any structural property, defined here as “complexity,” could associate with the network’s position in this space. We observed that the presence of two-component positive feedbacks is indeed one such property. Figure 2 shows how the number of these feedbacks (size of points normalized by median value of its population distribution, median value = 2) is enriched in networks with strong nonadditivity (the mean number of two-component positive feedbacks in nonadditive networks –those with ϵ_*T*_ beyond the 80th percentile of its population distribution– is 3.3. The expected value, based on 10,000 permutations, is 2.47, *p*-value = 0.005). This makes intuitive sense as these circuits act as switches of the dynamics in many biological situations [29]. Other measures of network complexity were also significantly higher for nonadditive networks (Methods and S2 Figure). These analyses support the intuition that more complex networks would lead to inferior PGS predictions.

### Genetic variation modulates predictability in a network-dependent manner

While we have seen how the mechanisms generating the phenotype influence the performance of the PGS, another important factor shaping its accuracy is the extent of genetic variation within the population [10, 16]. For instance, in a scenario with minimal variation in weights, it would be challenging to attribute phenotypic differences to them. We examined this aspect by contrasting situations with variation in all weights ranging from 5% (sd = 0.05) of their corresponding reference values to as much as 50% (sd =0.5; as compared to the 10%, sd = 0.1, used in previous sections). For each case, we compute the corresponding PGS. Variation affects prediction in a way dependent on the precise regulatory network.

Analyzing the same set of 100 networks, we identified various dependency patterns, including cases where specific ranges of variation result in either maximal or minimal prediction accuracy (Figure 3A; associated networks are detailed in S3 Figure). Patterns #1 and #2 generally correspond to networks that exhibit linear responses with low variation, leading to high prediction accuracy (*R*^2^). In contrast, patterns #3 and #4 represent nonlinear networks that increase their linear response and accuracy with greater genetic variance, but reach an optimal *R*^2^ for intermediate variation for pattern #3.

**Figure 3.**
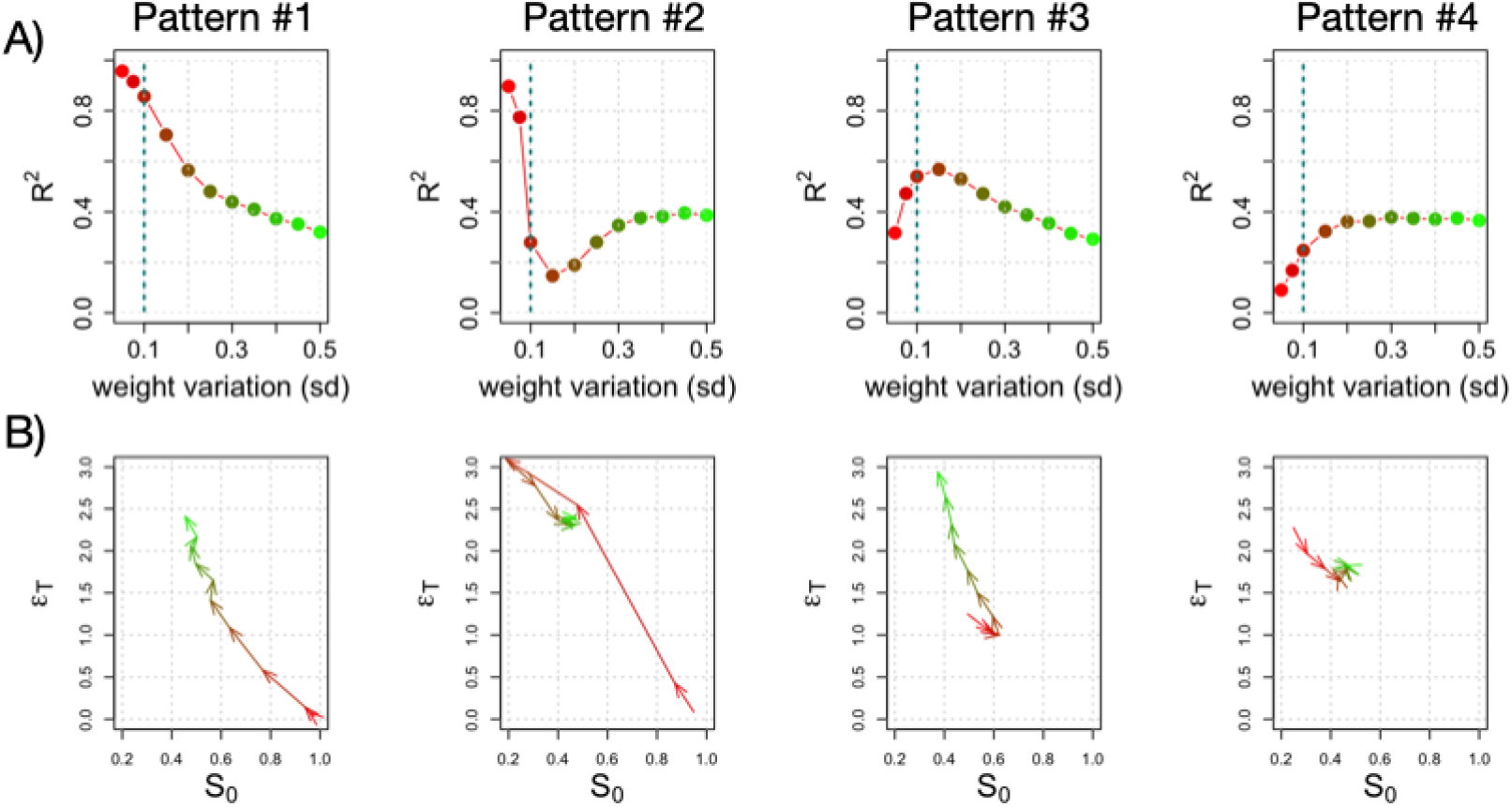
Genetic variation and its influence on prediction. **A)** We show four patterns of how PGS prediction accuracy (in terms of *R*^2^) changes as the genetic variation (sd of weights in the population) changes. The case of sd = 0.1 is indicated by dotted line. Patterns #1, #2 show strong prediction with small variation. Patterns #3, #4 show weak prediction with small variation that increases with more variation (#3 exhibits an optimal genetic variation for prediction). **B)** Trajectory of the patterns above in the additive space (*S*_0_, ϵ_*T*_). Red to green colors indicate a transition from lower to higher variation of the weights of the regulatory connections in the population.

This function can be understood using the additive space again, but this time we employ it to describe the impact of variation on each network (Figure 3B). Recall that the pair (*S*_0_, ϵ_*T*_) quantifies how much variation in all weights contributes additively or nonadditively to the phenotypic variation. Thus, pattern #1’s phenotypic variation is considerably explained by additive effects (large *S*_0_) in the range of small genetic variation, but this decreases as genetic variation increases (red to green in Figure 3, larger ϵ_*T*_), leading to a worsening of prediction accuracy. This is reflected in a diagonal trajectory towards the left in additive space. Other trajectories in this space also clarify the dependence on variation. For instance, the maximum in pattern #3 reflects a trajectory to the right (increased *S*_0_, decreased ϵ_*T*_) that returns to the regime of high nonadditivity and low additivity as variation intensifies.

Finally, it is important to evaluate how the additive or nonadditive impact of any weight on phenotypic variation shifts with changes in genetic variation. For a designated variation regime, we can obtain the Sobol indices (as shown in Fig.1C) and then compute the correlation between the indices associated with different regimes (S4 Figure). In some cases, the additive determinants change considerably depending on the variation within the population (e.g., #1 in S4 Figure), while in other cases, these determinants are relatively stable across conditions (e.g., #2 in S4 Figure). The nonadditivity of determinants typically increases with standard deviation but can sometimes remain relatively stable (e.g., #4 in S4 Figure). This example demonstrates that certain network architectures contribute to the stability of phenotypic determinants (see discussion). Overall, the effect of genetic variation on PGS accuracy is highly contingent on the network architecture responsible for generating the phenotype.

### Increased prediction accuracy is associated with the influence of core regulatory interactions

We can reinterpret the previous results in terms of the core/peripheral framework of the omnigenic model [18] by defining *core* interactions as those with their respective total Sobol index contributing to phenotypic variation beyond 1% 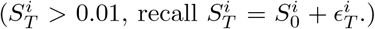 and the additive component bigger or equal than the nonadditive one 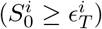, *peripheral* interactions as those with 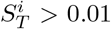, and 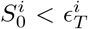 and *remote* interactions as those with 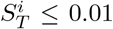. According to this, core and peripheral regulations are causally linked to the phenotype in an additive and nonadditive manner, respectively, while remote regulations exhibit a very loose connection.

Figure 4A shows that sorting the previous set of 100 networks by prediction accuracy reveals that as accuracy increases (larger *R*^2^ values), the proportion of core weight determinants increases, while the proportion of peripheral determinants decreases. Additionally, remote weights also exhibit an increase with *R*^2^ values. We also calculated the mean of the absolute effect size, |*β*|, associated with each determinant class (Figure 4B). Core determinants predominantly define the PGS beyond an accuracy threshold. In this manner, networks whose associated phenotypes are well predicted by an additive model would be responsive only to these core connections, which represent the essential determinants affecting the phenotype.

**Figure 4.**
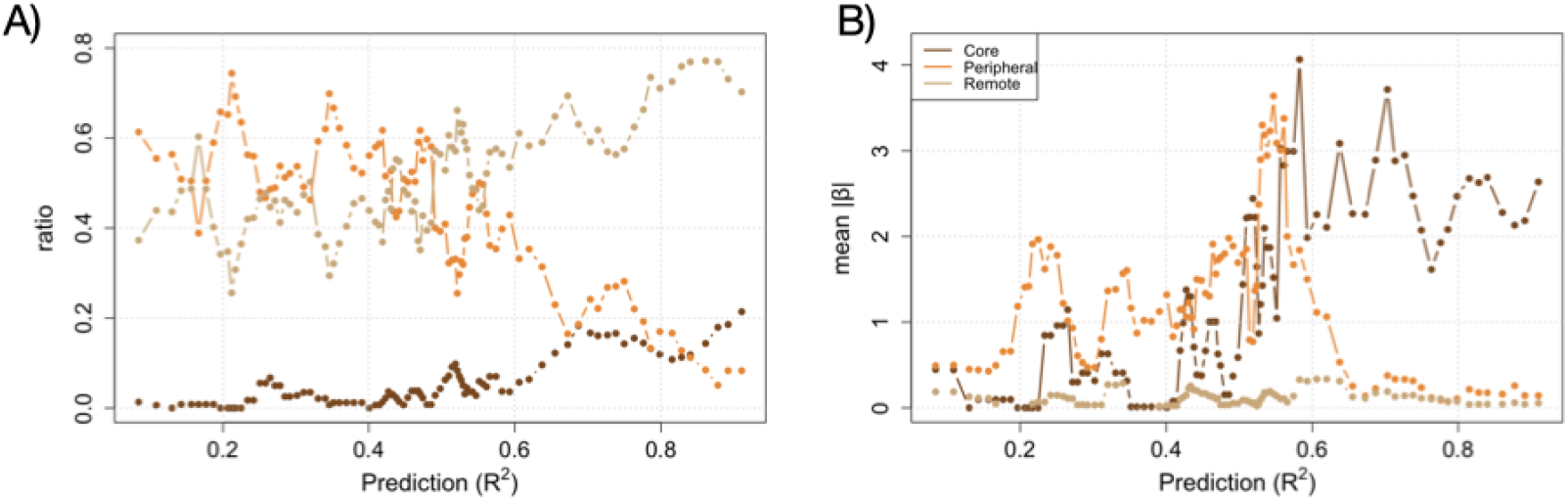
Core, peripheral, and remote determinants of the phenotype. **A)** Ratio of core, peripheral and remote regulatory connections over the total as function of prediction *R*^2^ for the set of 100 networks (with weight variation sd = 0.1; main text for definition of each connection class). Core determinants (dark brown line) increase with *R*^2^. **B)** Mean of PGS |*β*| ‘s as function of prediction. |*β*| ‘s at intermediate *R*^2^ are dominated by peripheral interactions (orange line). Both plots show rolling windows analysis of the relevant measure sorted by increasing *R*^2^, windows of size 5.

### PGSs show imperfect transferability

Finally, we investigated the challenge of applying a score developed under specific conditions to different ones. We examine two scenarios. We first employed a PGS computed with a given variation in the population to predict phenotypes of individual networks with weights sampled from a different distribution, a circumstance that holds significant implications in the explicit problem of the validity of GWAS findings across varied human population [24].

Figure 5A shows the *R*^2^ obtained when applying a PGS computed at sd = 0.1 to anticipate the phenotype of individuals whose weights are sampled from a different distribution (values normalized to the *R*^2^ obtained for individuals drawn for the original distribution, with sd=0.1). This is evaluated across the four patterns (networks) shown in Figure 3. We noticed that transferability tends to be effective for conditions comparable to the original training set and that it aligns with the patterns observed in Figure 3. Specifically, a PGS calculated with a population with sd of 0.1 demonstrates increased transferability in situations where it exhibits higher predictability. For instance, there is a noticeable rise in transferability for pattern #2, as depicted in the second row of Fig. 5A, within ranges where the sd is less than 0.1. This trend is consistent with the R^2^ values calculated for those corresponding regimes, as shown in Fig. 3A.

**Figure 5.**
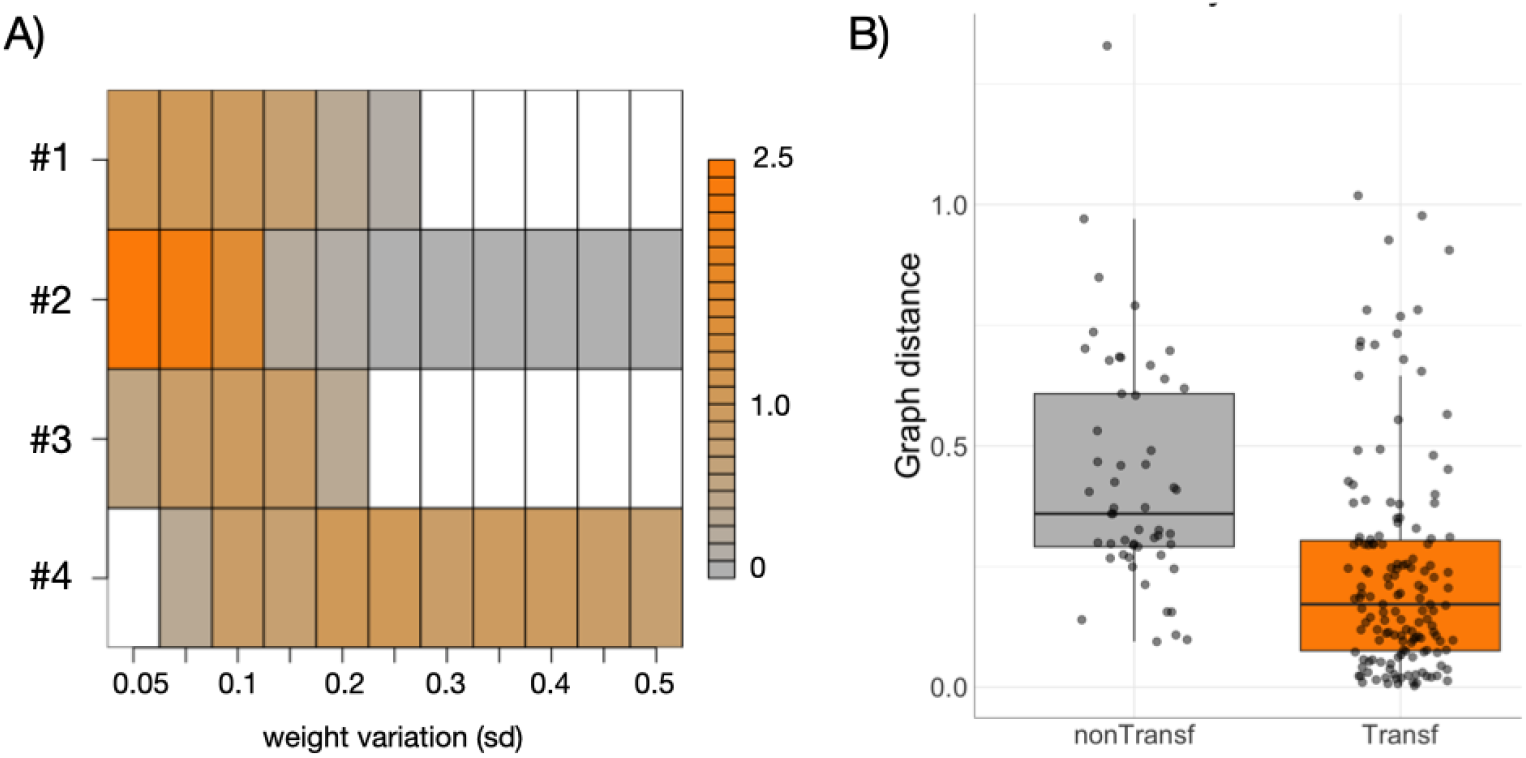
PGS transferability. Transferability of prediction to populations with different genetic variation, A), or network architecture, B). **A)** A PGS computed for each reference network in Figure 3, and a population weight variance of 10% (sd = 0.1), is applied to predict the phenotypes generated for each corresponding network under different population variations (weight variation). Each row is the corresponding *R*^2^ of the prediction normalized by the *R*^2^ of the reference condition (at sd=0.1). White denotes *R*^2^ < 0. **B)** A single regulatory connection of the network associated with pattern #1 (Figure 3) is randomly mutated, assigning a new weight with distribution ∼ 𝒩 (0, 1). The phenotypes generated with the mutant network are also predicted using the PGS computed with the “wild-type” network (with weight variance of sd = 0.1), and an R^2^ value is computed. Nontransferable mutant networks (*R*^2^ < 0) exhibit greater dissimilarity (to the wild-type) than transferable ones based on a spectral network (graph) distance (Wilcoxon rank sum test p-value < 0.001, Methods). Each dot represent a different mutant of a total of two hundred.

In the second situation, we first modified the reference network corresponding to pattern #1 (S3 Figure) by randomly changing one weight (Methods). We then asked if phenotypes generated with individual networks whose strengths vary over the new mutant network (sd = 0.1) can still be predicted using a PGS computed with the original reference network (also obtained with sd = 0.1). Using these two phenotypic datasets, we calculated an R^2^ as a measure of transferability. We additionally evaluated a dissimilarity score between the mutant and its corresponding reference network (network spectral distance, Methods). Repeating this protocol, we found that perturbations significantly altering the network were associated with low transferability (Figure 5B). These perturbations involved weights with typically large 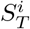 values (mean 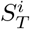 for nontransferable mutant graphs = 0.13, mean 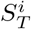 for transferable mutant graphs = 0.01, *p*-value < 10^−4^).

## Discussion

What are the limitations of additive statistical tools in capturing the complexities of genetic influences on complex traits? We approached this question by explicitly incorporating mechanistic gene regulation networks producing phenotypes. By introducing changes to the regulations, we can derive a combination of both genetic and phenotypic variations, allowing us to compute a PGS. In our scenario, the phenotype is directly associated with the steady state of the network.

The network framework is crucial as it offers a rationale for the multitude of determinants identified in GWAS studies [30]. These determinants are often categorized into a handful of core elements, characterized by variations that exert a substantial effect size on the phenotype. Conversely, numerous peripheral elements exhibit weaker effect sizes and are associated with regulatory effects that may not be directly linked to the specific trait under investigation. These are the constituents of the recently introduced omnigenic model [18, 19]; but see also [31].

We first use local stability analysis to study a specific example of PGS. This analytical framework captures the extent of linear behaviours in phenotype as regulatory strengths change, so we expected that it would shed partial light on causal variants of substantial effect size, as it did (Figure 1AB). But the approach is inherently limited to networks operating in a relatively linear dynamical regime. To better understand nonlinear cases, we applied global sensitivity techniques [26, 27]. We can then measure both the additive and nonadditive contributions of each regulatory connection to phenotypic variation. Our findings revealed that certain connections exhibit both additive and nonadditive effects, e.g., Figure 1C, thus reinforcing previous discussions regarding the impact of nonlinear genetic interactions, or biological epistasis, on additive genetic variance [14, 15, 32]. Furthermore, we observed that the nonlinearity and complexity of the network, assessed through stability analysis and metrics such as the presence of two-component positive feedback loops (Figure 2), correlate with its nonadditivity. Thus, we were able to partially link mechanistic and statistical features, a topic of ongoing debate [16]. Networks exhibiting greater linearity tend to yield higher prediction accuracy for the PGS, consistent with expectations.

### Broader implications

We utilize the sensitivity framework to address three critical topics within this context: the distribution of core, peripheral, and remote determinants, the impact of genetic variation on accuracy, and the transferability of the predictions. As illustrated in Figure 4, intermediate regimes of predictability exhibit a sparse distribution of core determinants and a multitude of peripheral/remote ones. This pattern closely mirrors what is observed in complex human traits [19]. For instance, the prediction accuracy of height, anatomical features being those with best predictions, is around *R*^2^ ∼ 0.2 [33, 34]. This would suggest that genomes are in a regime of low (statistical) prediction.

Moreover, an important component of genetic architecture that impacts the ability to predict a trait is genetic variation, defined by the distribution of alleles within a population [14]. While our modeling of variation incorporates only a normal distribution of regulatory weights around wild-type values, changes in the dispersion of this variation reveal how effectively an additive model can predict the phenotype. This effect is contingent upon the architecture of the network, i.e., the structure of the genotype-to-phenotype map [32, 35] (Figure 3). Beyond prediction accuracy, we observed a degree of stability in certain scenarios in the number and effect size of phenotypic determinants (both core and peripheral) across a range of genetic variation in the population (S4 Figure).

This last observation may help explain the similar distribution of selection coefficients at causal variants across traits, proposed as a reason for the consistency of their genetic architecture [36]. The authors suggested that such uniformity indicates an overlap in the numerous peripheral determinants contributing to trait variation. We reinforce this idea by emphasizing situations where phenotypic variation is associated with a relatively *fixed* set of causal factors, peripheral or not, in certain networks. This aspect, along with the possibility that gene regulation networks are consistent across different traits, would contribute to the prevalence of selection coefficients being correspondingly similar.

It has also been proposed that negative selection prevents the clustering of causal variants of the phenotype in a few core elements with significant effects, instead spreading them across a larger number of determinants with smaller effects, thereby “flattening” their distribution [37]. We propose that part of this flattening could also be related to the complexity of the networks, with networks acting in a more nonlinear way leading to a *de novo* effect size distribution spread across many determinants with small effects (S4 Figure, S5 Figure). Thus, beyond selective constraints, effect sizes and their distribution can also be influenced by network-level properties.

Lastly, regarding transferability, our initial findings demonstrate that PGSs exhibit a degree of transferability to populations with relatively similar genetic variation, as one might anticipate (Fig. 5A). Furthermore, we investigated how slight changes (single mutations) in the network architecture responsible for generating the phenotype could result in prediction failures (Fig. 5B). We observed that failures occur when changes predominantly involve mutations of *nonremote* connections 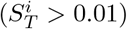. That phenotypes are only considerable affected by a subset of potential determinants echoes the concept of “sloppy” parameters; parameters that have little impact on the functioning of a biological system despite their change [38, 39]. Indeed, making use of this concept in our context, one could expect that combinations of nonremote weights would lead to stiff directions, where small changes result in significant alterations in network behavior and predictions.

### Final remarks and conclusions

Recent studies have also explored the limitations of additive models in capturing the impact of the genetic contribution to phenotypes, employing various mechanistic models that complement our approach. For example, Li et *al*. [1] investigated a deep neural network with varying layers to model the genotype to phenotype mapping. Similar to our findings, they observed that the degree of missing heritability, i.e., the fraction of phenotypic variance explained by genetic variance [10], is proportional to the complexity of the map (in their case, the number of layers in the neural network). In addition, we examined a framework based on genome-scale metabolic reconstructions to address the issue of predictability [40]. These models are more realistic as they encompass all known metabolic reactions and the genes encoding each enzyme for a given organism [41]. Despite the presence of nonadditive effects, we observed that the sum of additive terms accounts for over 75% of the total phenotype variation in this system. This additive component is contingent upon the structure of the metabolic genotype-phenotype map, particularly the monotonic relationship between gene content and phenotype [42, 43]. Moreover, much earlier work, such as that of Frank [44], highlighted how the structure of networks could delineate a stronger or weaker “zone of nonlinearity”, wherein changes in input have roughly linear effects on output. This observation resonates with our results, further corroborating the importance of network structure in delineating the boundaries of linearity in genotype-phenotype mappings.

Overall, our work helps assess the limitations of linear models that assume additive genetics. We employed a relatively simplistic framework, which still offers valuable insights into some of the current challenges associated with human PGSs. However, more realistic modeling is needed. Additionally, our findings underscore the plasticity of gene networks, which can shift their responses between linear and nonlinear functional regimes. Networks operating in a nonlinear regime might undergo *linearization* due to interactions with other networks (such as cell-cell interactions) or specific gene-environment interactions, potentially affecting prediction accuracy. The central challenge is to integrate various approaches to understand and predict complex phenotypes generated in such an intrinsically nonlinear manner [16] and to determine how these efforts can advance our predictions beyond merely “hoping for the best” [17].

## Methods

### Gene regulation network model

We consider regulatory reference networks constituted by N genes that regulate each other, where every possible regulation between genes is created with the same constant probability, *c* (0 ≤ *c* ≤ 1), according to an Erdös-Rényi model [45] (networks will not contain any self-loops) and stored in a matrix W. Weights on this matrix follow a normal distribution with mean 0 and standard deviation 1. The expression of the genes at a given time t is defined by a gene expression vector 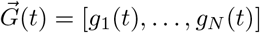 whose (discrete) time dynamics follows a set of nonlinear coupled difference equations: *g*_*i*_(*t* + 1) = *f*[∑*w*_*ij*_*g*_*j*_(*t*)]. Here, f is a sigmoidal function *f*(*x*) = 2/[1 + exp(−*ax*)] – 1, with *a* controlling the steepness of the sigmoid (the larger a, the steeper) that normalizes expression values in the range [-1 (`off` state of a gene), 1(on state of a gene)]. This model developed in [22] has its roots in statistical physics [46]. In our simulations, we considered *N* = 10, *c* = 0.3, and *a* = 10. Studies involving larger networks and alternative connectivities will be the focus of future work.

### Network dynamics

We set the initial state 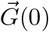 as a vector of *g*_*i*_(0)= 1 or -1, randomly assigned with probability 1/2. The states change following the difference equations defined before until it reaches an equilibrium. We selected genetic regulatory networks that lead to a fixed-point attractor equilibrium within a number of 100 iterations [22, 23]. We followed previous protocols to numerically identify when the equilibrium 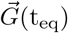 is reached. This is recognized when a measure of how much the state is changing for a specified time period is considerably small (*ψ* score from [23]). The number of iterations to equilibrium is named *path length*.

### Genetic and phenotypic variation

We generate genetic variation within a population by sampling gene-gene interaction weights from a normal distribution with mean equal to the specific weight *w*_*ij*_, of the corresponding reference network, and standard deviation given by 10% of that value, unless indicated otherwise. This distribution is relatively simple, incorporating only a basic variation around a wild-type value and avoiding the complexities of allele frequency distributions, e.g., [16]. Nevertheless, it represents an initial step toward incorporating variation into this modeling approach. Finally, for each member of a population, we maintained the same fixed initial state to compute the steady-state expression, the mean of which corresponds to the individual network phenotype. We considered a population size of 10^4^ individuals.

### Network nonlinearity

We quantify the nonlinear dynamical regime of a given reference network by computing the eigenvalues of its Jacobian at equilibrium [25]. We typically find a combination of eigenvalues equal to 0 and less than 1. When a discrete dynamical map as our case presents such combination, it suggests a mixture of stability conditions and dynamic behaviors. Eigenvalues equal to 0 indicate *neutral* stability, where the system remains at equilibrium without evolving over time. Perturbations around the equilibrium state do not cause the system to diverge or converge but rather stay in the vicinity of the equilibrium point. We computed the number of 0 eigenvalues for each network and showed that is associated with nonadditivity (S1 FigureB, right panel). Moreover, eigenvalues less than 1 indicate *stable* dynamics, where perturbations around the equilibrium state decay over time. Rapid decay –eigenvalues farther from 1 but still less than 1– suggests that the linearized model around that equilibrium is a good approximation. We showed that the maximal value of these eigenvalues for each reference network considered in Fig. 2 linked to additivity (S1 FigureB, left panel). Thus, this combination of zero eigenvalues and eigenvalues less than 1 defines a kind of mechanistic information that helps explain the statistical additive space.

### Network complexity

Other measures of network “complexity” beyond the presence of two-component feedbacks show significant differences, such as path length (how quickly the network reaches equilibrium gene expression; path length is larger the more nonadditive the network is; mean path length for nonadditive networks with ϵ_*T*_ beyond the 80th quartile, Fig. 2, is 17.9, expected value 15.96, *p*-value = 0.01), network density (the ratio of the number of edges to the number of possible edges) and clustering coefficient (the probability that adjacent nodes of a given node are connected). Both last scores are also significantly higher for nonadditive networks beyond a given ϵ_*T*_ threshold, defined just as previously (S2 Figure).

### Polygenic score

We define the network phenotype, y, as the *mean steady state expression* of its genes and use a high-dimensional regression framework for polygenic modeling and prediction: 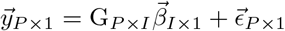, where *P* is the population size, *I* is the number of nonzero determinants, *y* is the vector of phenotypes, G is the genotype matrix, 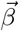 is the vector of effect sizes of the genes, and 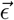 is some noise assumed normal with unknown variance. The components of the G matrix, which are the weights, are continuous in this case. This is analogous to how single nucleotide polymorphism genotypes are represented as continuous values after imputation [8].

The generated data was fitted using the Least Absolute Shrinkage and Selection Operator (`LASSO`), a type of regression that, within Bayesian statistics, assumes prior Laplace distributions for each coefficient rather than uniform distributions, as in the case of Ordinary Least Squares. As a result, with `LASSO`, some parameters are automatically shrunk to zero [47] making it a remarkable alternative to pruning and thresholding (P+T) or other regularization methods [6, 48]. Additionally, we determine the optimal value of the shrinkage parameter using five-fold cross-validation.

### Numerical computation of sensitivity indices

We followed [27]. In brief, we first generate a (*N*_sobol_, 2*k*) matrix of random numbers (k is the number of regulatory connections) and define two matrices of data (A and B), each containing half of the sample. *N*_sobol_ was taken to be 5 × 10^4^. We define a matrix C_*i*_ formed by all columns of A except the *i*-th column, which is taken from B. We then compute the model output for all the input values across the sample matrices A, B, and C_*i*_, obtaining three vectors of model outputs of dimension 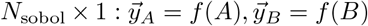 and 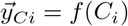. From these vectors we can compute the first- and total-effect indices for a given weight [27]; see also supplement in [40]. The bars in Fig. 1 represent the mean values of these indices, with small standard deviations (not shown for simplicity of presentation).

### PGS transferability

To assess transferability, we obtain a PGS in a reference condition and then apply it to predict phenotypes under various conditions, such as different standard deviations or networks. We calculate the corresponding *R*^2^ between the predicted and observed phenotypes as a measure of transferability. The graph spectral distance between two networks serves as a measure of dissimilarity between them. Spectral distance is defined as 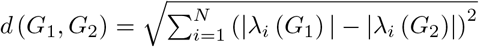, where |λ_1_ (*G*|) > |λ_2_ (*G*) |, … are the absolute value of the eigenvalues of each network in decreasing order and N the number of nodes.

## Supporting

### S1 Figure

**State flip and network nonlinearity. A)** We defined a “state flip” as occurring when a displacement *δw* from a wild-type weight value w generates a change in the sign of the expression of a given gene *i* from time t to time *t* + 1, i.e., sign[*g*_*i*_(t)] sign[*g*_*i*_(t + 1)]. Recall that *g*_*i*_(*t* + 1) = *f*[*w*_*ij*_*g*_*j*_(*t*)], with *f* being a sigmoidal function *f*(*x*) = 2/[1 + exp(−*ax*)] – 1. This flip is an indication of the nonlinear dynamics. **B)** We assess the nonlinear dynamics of each reference network in Fig. 2 based on the maximal absolute value of the eigenvalues of the Jacobian matrix at equilibrium (left panel) and the count of eigenvalues with a value of zero (right panel). A smaller dot on the left indicates a lower value of this eigenvalue, reflecting higher stability of the network. Networks with strong additivity—those with *S*_0_ beyond the 70th percentile of the population distribution—exhibited smaller values for these maximal nonzero eigenvalues (mean observed = 0.1, mean expected by random permutation = 0.17, *p* < 0.03, after 10,000 permutations). Conversely, a larger dot on the right indicates a higher count of zero eigenvalues, also signifying greater stability. Networks with strong nonadditivity—those with ϵ_*T*_ beyond the 70th percentile of the population distribution—tended to have fewer zero eigenvalues (mean observed = 0.47, mean expected by random permutation = 0.82, *p* < 0.02, after 10,000 permutations). For more details, see the network nonlinearity subsection in the main text and Methods.

### S2 Figure

**Network nonlinearity and prediction accuracy. A)** The ratio of the number of edges to the total number of edges (network density) is significantly bigger in networks with ϵ_*T*_ beyond the 80th quartile of the population distribution that are associated to lower prediction accuracy (*R*^2^). **B)** The probability that adjacent nodes of a given node are connected (clustering coefficient) is significantly bigger in nonadditive networks with ϵ_*T*_ beyond the 80th quartile. Histograms show the null distribution of either score (10000 randomizations of the assignment of ϵ_*T*_ values to the related score) and red vertical dashed line the corresponding observed value. See also Figure 2, main text.

### S3 Figure

**Network structures**. Networks corresponding to patterns #1 to #4, A) to D), respectively, in Figure 3. Associated R^2^ at sd= 0.01 are 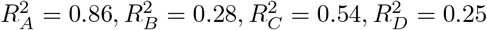.

### S4 Figure

**Additive and nonadditive effects and genetic variation**. For each pattern in Figure 3 (A-D) (networks shown in Fig. S3) we computed the correlation between the additive, 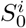 or nonadditive, 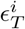, contributions to variance computed for different genetic variation (top figures on each panel). We also computed how these contributions change for each weight determinant (bottom figures on each panel). This plot highlights the range of stability of the additive/non additive role of each weight on the phenotypic variance. Note that some cases, e.g., #4, show considerable stability.

### S5 Figure

**Polygenicity, effect size and nonadditivity**. Polygenicity quantified as the number of weights with effect size *β* different from zero tends to decrease with mean effect size and increase with nonadditivity, although the tendency is weak but illustrative. Figures show the values for each of the 100 random network and a linear regression (continuous line) and confidence intervals (filled points); A) *R*^2^ = 0.046, B) *R*^2^ = 0.013.

## Data availability

Data and code for this work is available at Zenodo https://zenodo.org/records/13359410

## Acknowledgments

Thanks to all members and visitors of the LoGS lab for the discussions. This work was supported by grant PID2019-106116RB-I00 from the Spanish Ministry of Economy and Competitiveness with European Regional Development Fund.

