## Supplement for "Polygenic prediction and gene regulation networks"

Juan F Poyatos

**Logic of Genomic Systems Laboratory (CNB-CSIC)**
**28049 Madrid, Spain**


### Supplementary Figures

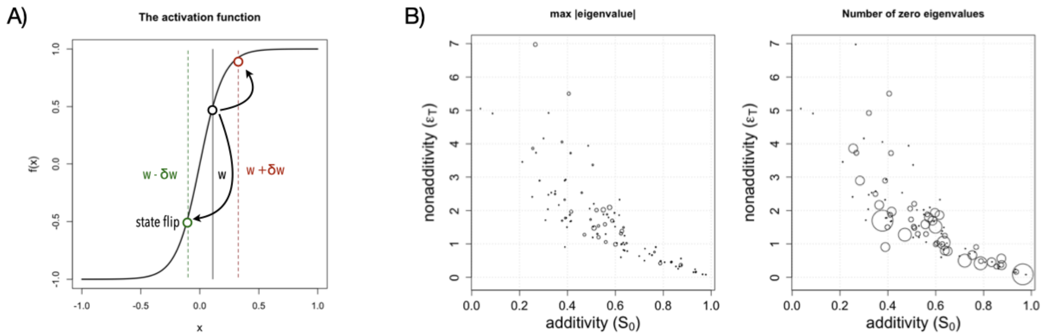


**Figure S1. State flip and network nonlinearity. A)** We defined a "state flip" as occurring when a displacement $\delta w$ from a wild-type weight value $w$ generates a change in the sign of the expression of a given gene $i$ from time $t$ to time $t+1$, i.e., $\text{sign}[g_i(t)] \neq \text{sign}[g_i(t+1)]$. Recall that $g_i(t+1) = f[\sum w_{ij} g_j(t)]$, with $f$ being a sigmoidal function $f(x) = 2/[1 + \exp(-ax)] - 1$. This flip is an indication of the nonlinear dynamics. **B)** We assess the nonlinear dynamics of each reference network in Fig. 2 based on the maximal absolute value of the eigenvalues of the Jacobian matrix at equilibrium (left panel) and the count of eigenvalues with a value of zero (right panel). A smaller dot on the left indicates a lower value of this eigenvalue, reflecting higher stability of the network. Networks with strong additivity—those with $S_0$ beyond the 70th percentile of the population distribution—exhibited smaller values for these maximal nonzero eigenvalues (mean observed = 0.1, mean expected by random permutation = 0.17, $p < 0.03$, after 10,000 permutations). Conversely, a larger dot on the right indicates a higher count of zero eigenvalues, also signifying greater stability. Networks with strong nonadditivity—those with $\epsilon_T$ beyond the 70th percentile of the population distribution—tended to have fewer zero eigenvalues (mean observed = 0.47, mean expected by random permutation = 0.82, $p < 0.02$, after 10,000 permutations). For more details, see the network nonlinearity subsection in the main text and Methods.

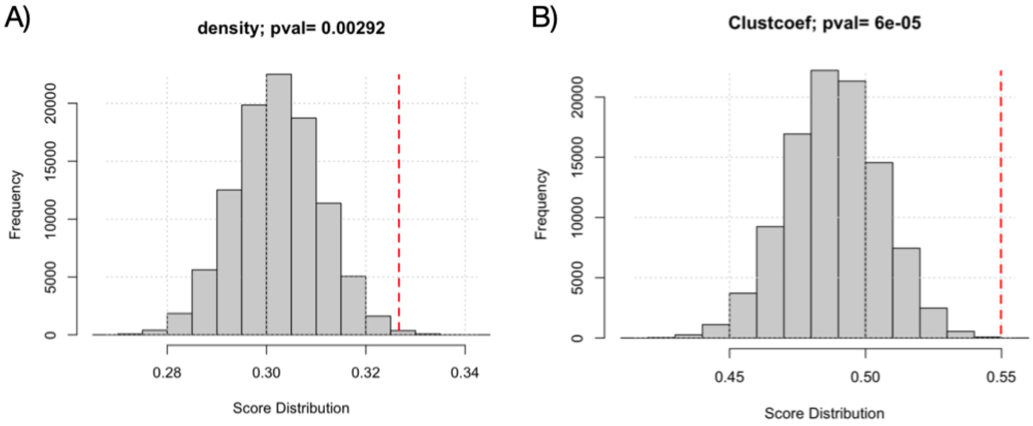


**Figure S2. Network nonlinearity and prediction accuracy. A)** The ratio of the number of edges to the total number of edges (network density) is significantly bigger in networks with $\epsilon_T$ beyond the 80th quartile of the population distribution that are associated to lower prediction accuracy ($R^2$). **B)** The probability that adjacent nodes of a given node are connected (clustering coefficient) is significantly bigger in nonadditive networks with $\epsilon_T$ beyond the 80th quartile. Histograms show the null distribution of either score (10000 randomizations of the assignment of $\epsilon_T$ values to the related score) and red vertical dashed line the corresponding observed value. See also Figure 2, main text.

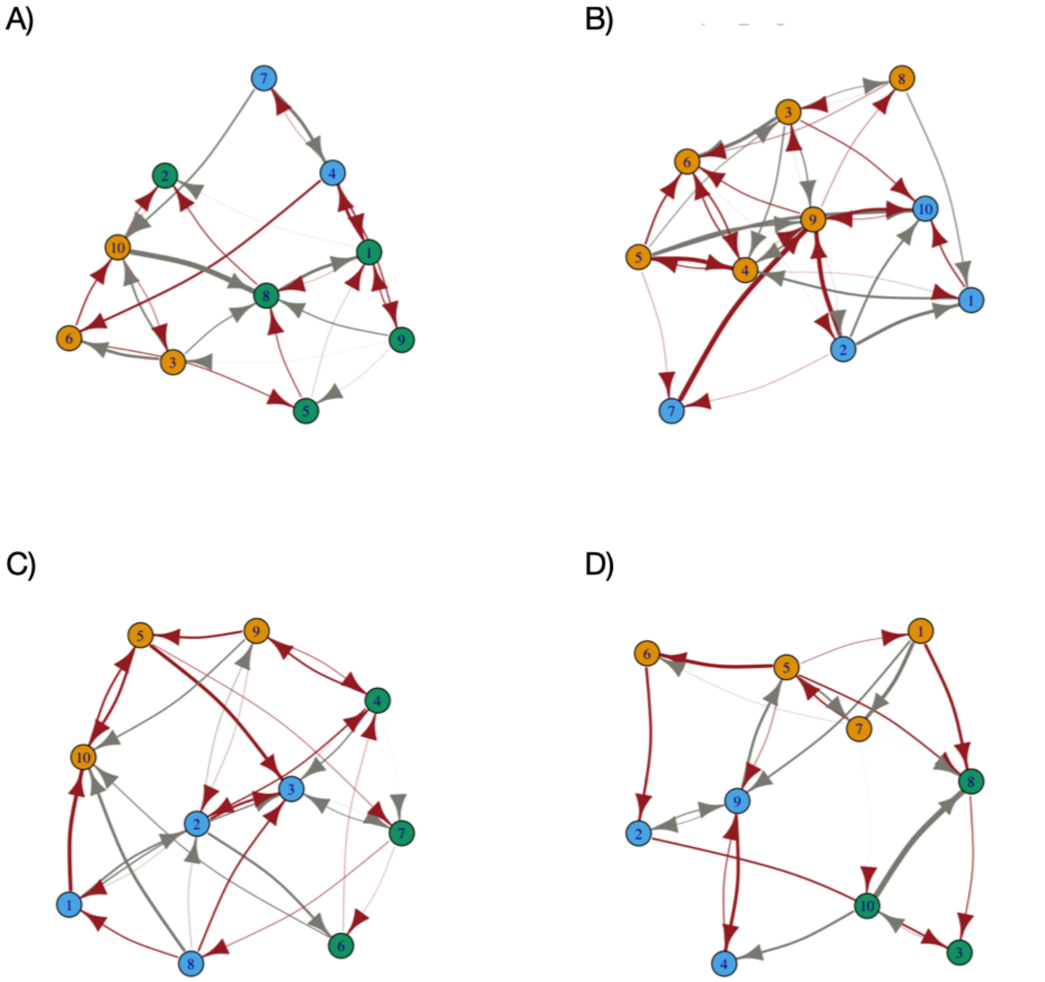


**Figure S3. Network structures**. Networks corresponding to patterns #1 to #4, A) to D), respectively, in Figure 3. Associated $R^2$ at sd= 0.01 are $R^2_A = 0.86, R^2_B = 0.28, R^2_C = 0.54, R^2_D = 0.25$.

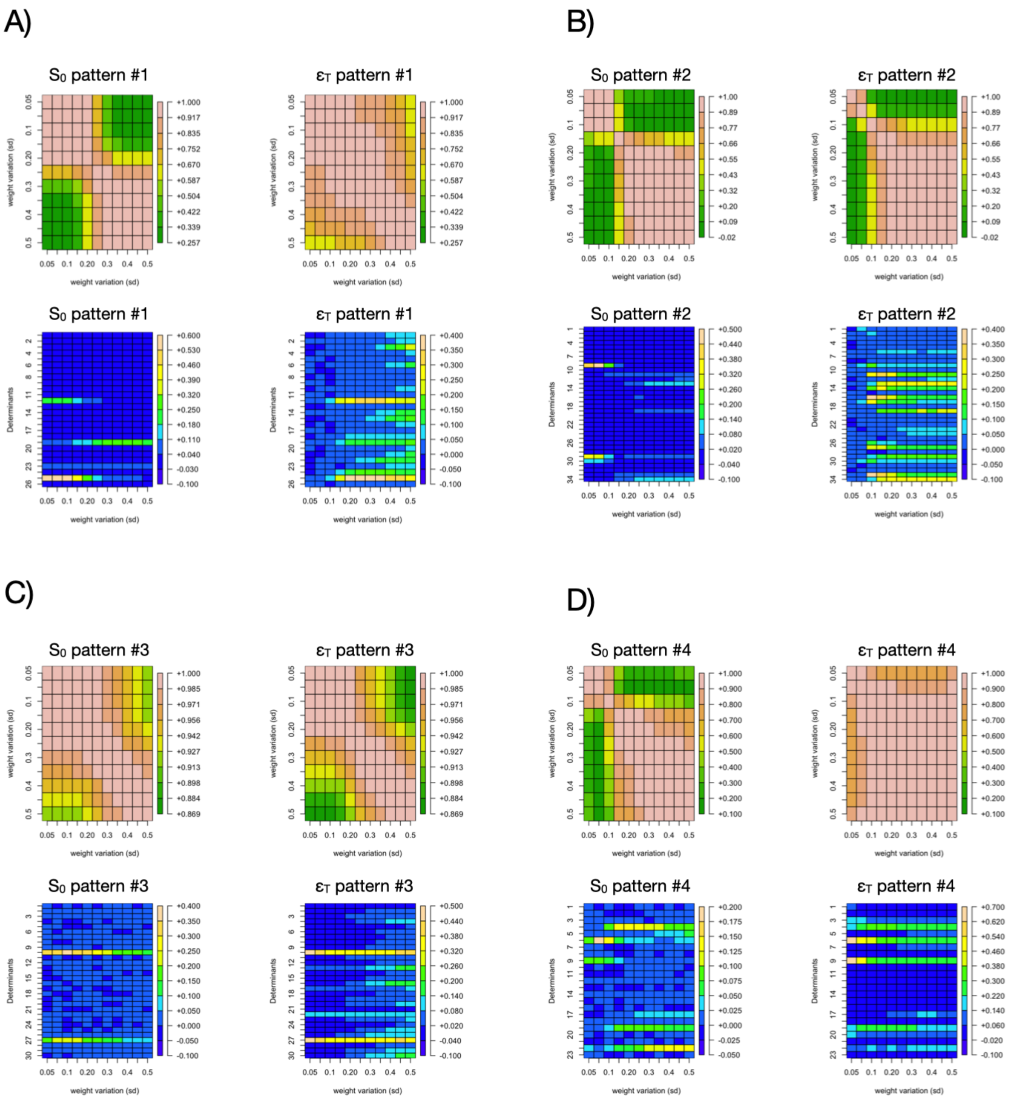


**Figure S4. Additive and nonadditive effects and genetic variation.** For each pattern in Figure 3 (A-D) (networks shown in Fig. S3) we computed the correlation between the additive, $S_0^i$ or nonadditive, $\epsilon_T^i$, contributions to variance computed for different genetic variation (top figures on each panel). We also computed how these contributions change for each weight determinant (bottom figures on each panel). This plot highlights the range of stability of the additive/non additive role of each weight on the phenotypic variance. Note that some cases, e.g., #4, show considerable stability.

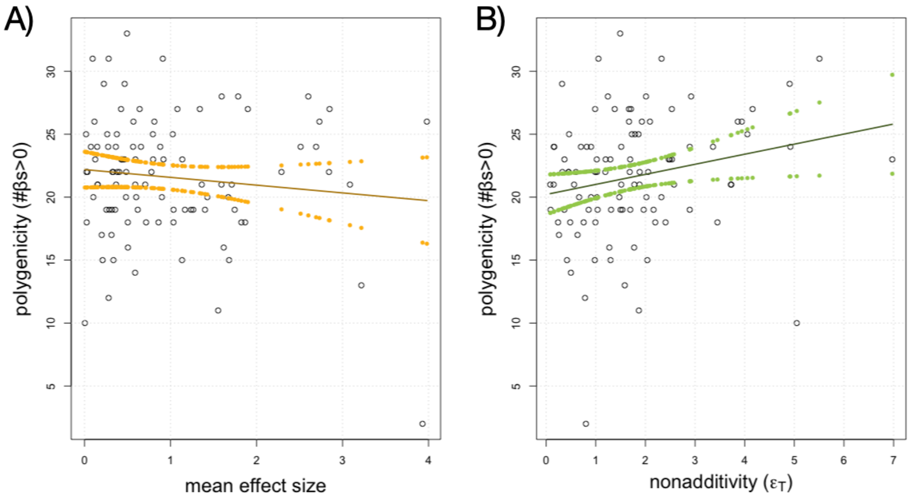


**Figure S5. Polygenicity, effect size and nonadditivity.** Polygenicity quantified as the number of weights with effect size $\beta$ different from zero tends to decrease with mean effect size and increase with nonadditivity, although the tendency is weak but illustrative. Figures show the values for each of the 100 random network and a linear regression (continuous line) and confidence intervals (filled points); A) $R^2 = 0.046$, B) $R^2 = 0.013$.
